# Kevlar: a mapping-free framework for accurate discovery of *de novo* variants

**DOI:** 10.1101/549154

**Authors:** Daniel S. Standage, C. Titus Brown, Fereydoun Hormozdiari

**Affiliations:** Population Health and Reproduction, University of California, Davis; Genome Center, University of California, Davis; MIND Institute, University of California, Davis; Biochemistry and Molecular Medicine, University of California, Davis; 1 Shields Ave, Davis, CA 95616

**Keywords:** *k*-mer, variant discovery, *de novo* variation, indel, autism, human genomics

## Abstract

**Motivation:** Discovery of genetic variants by whole genome sequencing has proven a powerful approach to study the etiology of complex genetic disorders. Elucidation of all variants is a necessary step in identifying causative variants and disease genes. In particular, there is an increased interest in detection of *de novo* variation and investigation of its role in various disorders. State-of-the-art methods for variant discovery rely on mapping reads from each individual to a reference genome and predicting variants from difference observed between the mapped reads and the reference genome. This process typically results in millions of variant predictions, most of which are inherited and irrelevant to the phenotype of interest. To distinguish between inherited variation and novel variation resulting from *de novo* germline mutation, whole-genome sequencing of close relatives (especially parents and siblings) is commonly used. However, standard mapping-based approaches tend to have a high false-discovery rate for *de novo* variant prediction, which in many cases arises from problems with read mapping. This is a particular challenge in predicting *de novo* indels and structural variants.

**Results:** We have developed a mapping-free method, Kevlar, for *de novo* variant discovery based on direct comparison of sequence content between related individuals. Kevlar identifies high-abundance *k*-mers unique to the individual of interest and retrieves the reads containing these *k*-mers. These reads are easily partitioned into disjoint sets by shared *k*-mer content for subsequent locus-by-locus processing and variant calling. Kevlar also utilizes a novel probabilistic approach to score and rank the variant predictions to identify the most likely *de novo* variants. We evaluated Kevlar on simulated and real pedigrees, and demonstrate its ability to detect both *de novo* SNVs and indels with high sensitivity and specificity.

**Availability:** https://github.com/kevlar-dev/kevlar

## Introduction

It is speculated that genetic variation is a major contributing factor in complex genetic disorders. The genetic heritability of many disorders is estimated to be relatively high. For example, the heritability of autism spectrum disorder (ASD) is over 0.6, and the heritability of Schizophrenia is over 0.5.^1, 2^ Only a fraction of this heritability is explained by known genetic variants, however, a phenomenon termed “missing heritability”.^3^ One hypothesis is that *de novo* mutations, in particular indels and structural variants, are a large source of causative variation (and consequently missing heritability) in developmental disorders.^3–5^ However, the complexity of *de novo* variant discovery, especially *de novo* indel and structural variant (SV) discovery, has resulted in an incomplete accounting of their contribution to these disorders. The discovery of genetic variants in general, and *de novo* variants in particular, remains a topic of intense research interest. In addition to illuminating the role of genetic variation in the etiology of complex disorders, improved discovery and cataloguing of *de novo* variants across many samples or cohorts will shed additional light on important unresolved questions in human genomics, including rates, biases, and mechanisms of new mutation.

Whole genome sequencing of simplex families (presenting an isolated case of a genetic disorder) is a proven successful approach for discovery of novel genetic variants resulting from *de novo* mutation in the germline.^5–9^ A “trio” comprised of an individual affected by the disorder (the proband), the mother, and the father (alternatively, a “quad” or “quartet” comprised of the proband, mother, father and a sibling) provides a rich information source for discriminating between shared and unique variation. Following standard variant calling protocols, mapping-based methods for *de novo* variant prediction begin by aligning reads to the reference genome. Variants are then predicted for each individual based on artifacts observed in the read alignments, such as mismatches, gaps, abrupt shifts in coverage, and discordant read pair distances or orientations.^10–16^ This initial process typically results in millions of variant predictions, which *de novo* variant discovery algorithms must examine to discern between inherited variation, true *de novo* variation, and spurious variant calls.

While reference-based variant discovery methods have proven valuable in the study of complex genetic disorders, we note some of their limitations. Despite consistent improvements in read alignment algorithms, finding the correct mapping for each read is still complicated by sequencing errors, repetitive DNA content, and mis-assemblies in the reference. Reads that do not map to the reference genome because they span mutation breakpoints or contain novel sequence are ignored completely by mapping-based variant predictors. Also, few methods are able to predict multiple variant types simultaneously using a single strategy, instead focusing exclusively on SNVs, short indels, or SVs separately. Finally, most variant calls determined by analysis of read alignments are not unique to the individual of interest (child, or *proband*) but instead reflect divergence in ancestry between the family and the reference genome donors. Estimates of human germline mutation rates give an expectation of approximately 80 novel mutations per generation,^17^ and distinguishing true *de novo* variation events from millions of inherited or false variants is a substantial challenge.

More generally, accurate and comprehensive *de novo* variant discovery is complicated by several computational and biological factors, and remains an elusive goal. Any algorithm must be confident not only in the *existence* of the variant in the proband but also its *non-existence* in both parents. And while single nucleotide variants (SNVs) are the most common variant type, larger variants that are less frequent nevertheless affect more nucleotides overall and are hypothesized to have an even greater impact in genetic disorders. Accurate discovery of these larger *de novo* variants is particularly challenging due to the inherent complexity of indel and SV prediction. In a reference-mapping context, calling indels with confidence requires accurate mapping of each read spanning the indel, with all gaps arranged consistently. This is possible only for short indels, and tends to be prone to error and misalignment. Thus, prediction of indels with length >10 bp has proven to be very challenging and accompanied by high false positive and false negative rates. Furthermore, the prediction of SVs via read mapping is only possible through indirect signatures such as alterations in read depth or read-pair signatures. These signatures can be quite noisy and result in high rate of false negative and false positive prediction. As a result, some basic properties of *de novo* SVs, including their rate of occurrence, remain unknown. It is important to note that there also exists no method for predicting more complex types of *de novo* SVs, such as inversion-duplication.

Many of the challenges with *de novo* variant prediction can be mitigated by an approach that compares sequence content between related individuals directly, rather than indirectly via a reference genome. Such an approach does not require any read alignments, nor is it sensitive to off-target shared/inherited variation. What a mapping-free approach *does* require is a signature of variation that is not defined in terms of artifacts observed in read alignments.

One of the first tools to explore a mapping-free strategy for predicting and genotyping variants was Cortex, which introduced the concept of a “colored de Bruijn graph” to compare sequence content from two or more samples and predict variants between samples.^18^ Cortex was used successfully for predicting variants in the 1000 Genomes Project. The DISCOSNP method^19^ implemented a very efficient strategy for scanning a de Bruijn graph for “bubbles” reflective of isolated SNVs. More recently, DiscoSnp++ has improved on this strategy and is capable of predicting isolated SNVs, proximal SNVs, and indels without the use of a reference genome.^20^ At the core of both methods is the analysis of *k*-mers, or sequences of a fixed length *k*.

Increased attention is being given to these kinds of *k*-mer based methods that avoid read alignments altogether. Indeed, mapping-free strategies for a variety of genomic and transcriptomic applications have become increasingly prominent, in large part due to their efficiency and robustness to the shortcomings of reference genomes. (It is important to note that these and other developments have greatly benefited from the availability of software libraries for rapid exact and approximate *k*-mers; These libraries include Jellyfish,^21^ khmer,^22^ ntHash,^23^ DSK,^24^ and KMC.^25^) In the realm of transcriptome analysis, tools such as Kallisto^26^ and Sailfish^27^ are capable of accurate RNA-Seq quantification at a fraction of the time and computational cost of previous mapping-based strategies. A recent study has also introduced a novel mapping-free method for performing genome wide association studies (GWAS) from whole genome sequence data^28^ using *k*-mer counts. The tool HAWK^28^ performs rapid and accurate discovery of variant-phenotype associations by directly comparing *k*-mer frequencies between arbitrary numbers of case and control samples. HAWK counts all *k*-mers in the sequenced samples and finds *k*-mers which are significantly associated with the phenotype/trait of interest (“significant *k*-mers”), and then performs a local assembly of these significant *k*-mers to predict the corresponding significant variants associated with the traits. This approach provides an efficient method for discovery of significant associations between all types of variants (i.e., SNVs, indels, and SVs) and the phenotype/trait of interest.^28^

Developments in variant prediction frameworks continue to spur improvements in a variety of contexts. Scalpel^29^ implements a hybrid method for *de novo* indel discovery from whole-exome sequencing of quads. Read mapping is used only to localize reads to the reference genome. In subsequent steps, Scalpel performs localized *de novo* assembly of reads at loci of interest and aligns assembled contigs back to the loci to annotate any *de novo* variants present.^29^ More recently, NovoBreak^30^ introduced a method that utilizes *k*-mer counts to predict somatic variants, including structural variants, by comparison of paired tumor and normal whole-genome sequence samples. COBASI^31^ performs rapid and accurate *de novo* SNV discovery on whole-genome sequencing of trios by computing perfect matches to unique strings in the reference genome and then identifying abrupt shifts in the coverage of the resulting alignments Finally, mapping-free approaches such as LAVA,^32^ VarGeno,^33^ MALVA^34^ and Nebula^35^ were recently developed for fast and accurate genotyping of common variation using whole-genome sequencing data.

The present study introduces a new mapping-free strategy grounded on a *k*-mer based formulation of the *de novo* variant discovery problem. Intuitively, a novel germline mutation should result in new sequence content in the proband compared to the parental genomes. Even in the simplest case, a single nucleotide substitution, most of the *k*-mers spanning the mutation should be unique, given a sufficiently large value of *k*. Incidentally, this is also true for other classes of variants, such as indels and various types of structural variation. And with sufficiently deep sampling of the proband genome, the expectation is that these novel *k*-mers are present in the read data at levels that can be readily distinguished from sequencing errors. Thus, it should be possible to detect both single nucleotide variants and larger variants (indels, SVs) simultaneously using a single mapping-free model.

Building on this intuition, we developed Kevlar, a new method based on a mapping-free formulation of the *de novo* variant discovery problem. Kevlar examines *k*-mer abundances to identify “interesting” *k*-mers, which we define as having significantly high abundance in the proband/child reads while being effectively absent in the reads from both parents. These interesting *k*-mers are an indicator of the potential existence of a *de novo* variant in the proband, and are conceptually similar to the “significant” *k*-mers used by HAWK.^28^ We next group the reads containing interesting *k*-mers into disjoint sets, each reflecting a putative variant, based on the *k*-mers shared between pairs of reads. Kevlar then uses standard algorithms to assemble each set of reads into contigs and align the assembled contigs to a reference genome to make preliminary variant calls. Finally, Kevlar employs a probabilistic model to score predicted variants to distinguish between miscalled inherited variants and true *de novo* mutations.

We demonstrate the utility of this new method on simulated and real data. We show that Kevlar achieves similar predictive performance to best-in-class tools for SNV and short indel discovery, while at the same time predicting larger events with high accuracy. We also demonstrate Kevlar’s ability to accurately predict large-scale SV events, defining breakpoints with nucleotide-level precision.

Kevlar is available as an open source software project, and can be invoked via a Python API, a command-line interface, or a standard Snakemake workflow.^36^ The stable and actively developed source code is available at https://github.com/kevlar-dev/kevlar, and documentation is available at https://kevlar.readthedocs.io.

## Results

We present a novel framework for discovery of *de novo* variants based on direct comparisons of sequence content between related individuals, requiring no mapping of short reads to a reference genome. This framework utilizes a single strategy that accurately predicts single nucleotide variants (SNVs), insertions and deletions (indels), and structural variation events simultaneously.

### Overview of Kevlar

Our variant discovery strategy is fundamentally a search for novel DNA content in the sample of interest. It is based on the intuition that a *de novo* mutation will result in novel sequence that can be detected by analysis of *k*-mers, or short subsequences of length *k*. Often the subject is a child affected by a disorder or other trait of interest (referred to as *proband*), with related individuals being the two parents.

Figure 1d summarizes the Kevlar workflow. In brief, DNA sequence reads from the case and control samples are processed independently. For each sample, the reads are split into overlapping subsequences of length *k*, or *k*-mers, and the abundance of each *k*-mer is stored for subsequent lookup. A second pass over reads from the case sample then identifies all *k*-mers that are unique to the proband—that is, *k*-mers that are abundant in the proband but effectively absent in both parents. Reads containing any novel *k*-mers are retained for subsequent processing.

**Fig 1 a.**
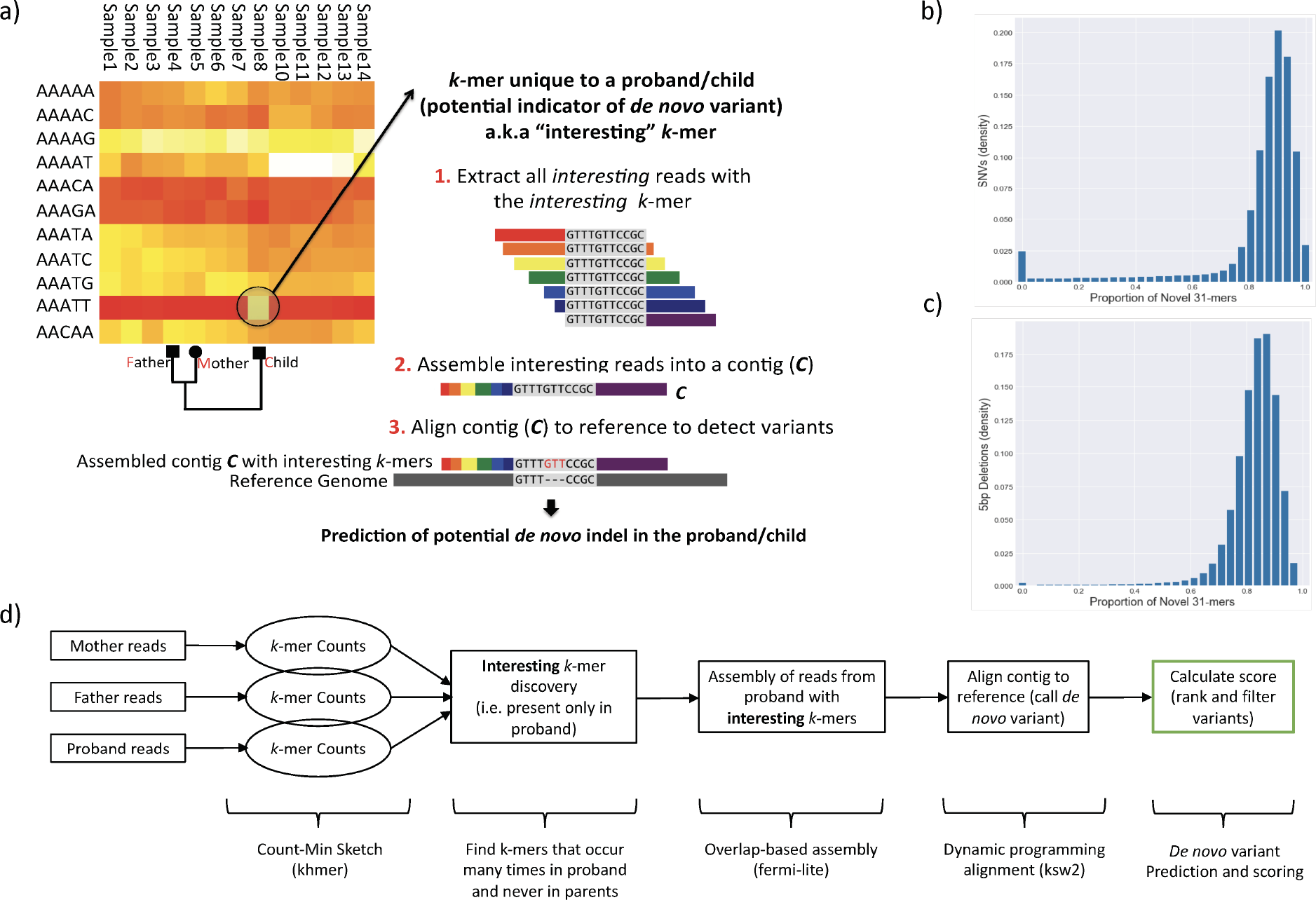
The overall mapping-free approach for *de novo* variant discovery b) The proportion *N*_*m*_ of *k*-mers associated with a simulated SNV mutations that are novel, aggregated over all sites in the genome. The trend observed for *k* = 31 holds for a wide range of *k* values (≈ 20− ≈ 60) c) The proportion of *k*-mers associated with a simulated 5bp indels that are novel, aggregated over all sites in the genome. d) The Kevlar pipeline․.

After applying filters for contamination and erroneous *k*-mer abundances, the reads containing novel *k*-mers are partitioned such that any two reads sharing at least one novel *k*-mer are grouped together. The reads in each partition are then analyzed independently: they are assembled into a contig, the contig is aligned to the reference genome, and the alignment is used to assess the presence of a variant and make a variant call. Finally, Kevlar employs a likelihood based score to rank and filter the variant calls.

Each step of the Kevlar workflow is discussed in detail in the Materials and Methods section.

### Performance on simulated data

We simulated whole-genome shotgun sequencing of a hypothetical (constructed) family for a fine-grained assessment of Kevlar’s accuracy in identifying different variant types at different levels of sequencing depth. Our simulation realistically modeled the inheritance of parental variants, but also included hundreds of “*de novo*” (unique to the proband) SNVs and indels ranging in size from <10bp to 400bp. The sequencing was simulated at 10x, 20x, 30x, and 50x coverage with low error rate. We compared Kevlar’s accuracy on this data set to two widely used mapping-based *de novo* variant callers (GATK PhaseByTransmission^37^ and TrioDenovo^38^) as well as two mapping-free/hybrid variant callers (Scalpel^29^ and DiscoSnp++^20^).

The accuracy of all variant callers evaluated is poor at low (10x) coverage (see Figure A1). GATK PhaseByTransmission makes very few variant predictions at 10x coverage. The remaining variant callers report numerous predictions, but in general suffer from both low sensitivity (failing to predict many true variants) and poor specificity (predicting many false variants). Triodenovo shows the best prediction performance for SNVs and short (1-100bp) indels at 10x coverage. At 20x coverage (Figure A2), all five algorithms show marked improvement in SNV detection, in particular TrioDenovo which achieves ≥90% sensitivity. Scalpel exhibits both improved sensitivity and improved specificity at 20x coverage, and approaches or surpasses Triodenovo’s performance for indels of most lengths. Kevlar’s ability to accurately detect indels > 100 bp becomes evident at 20x coverage.

At higher levels of coverage (30x and 50x), Kevlar consistently achieves top performance across all variant types (see Figures 2 and A3). Notably, Kevlar recovers ≥90% of true variants while making very few false predictions across all variant types at high coverage. TrioDenovo shows marginally better sensitivity than Kevlar for predicting SNVs at 30x and 50x (as does GATK PhaseByTransmission at 50x), but at the expense of numerous false predictions. Kevlar also rivals Scalpel’s impressive short indel prediction performance, and exceeds it for predicting long (> 100 bp) indels.

**Fig 2.**
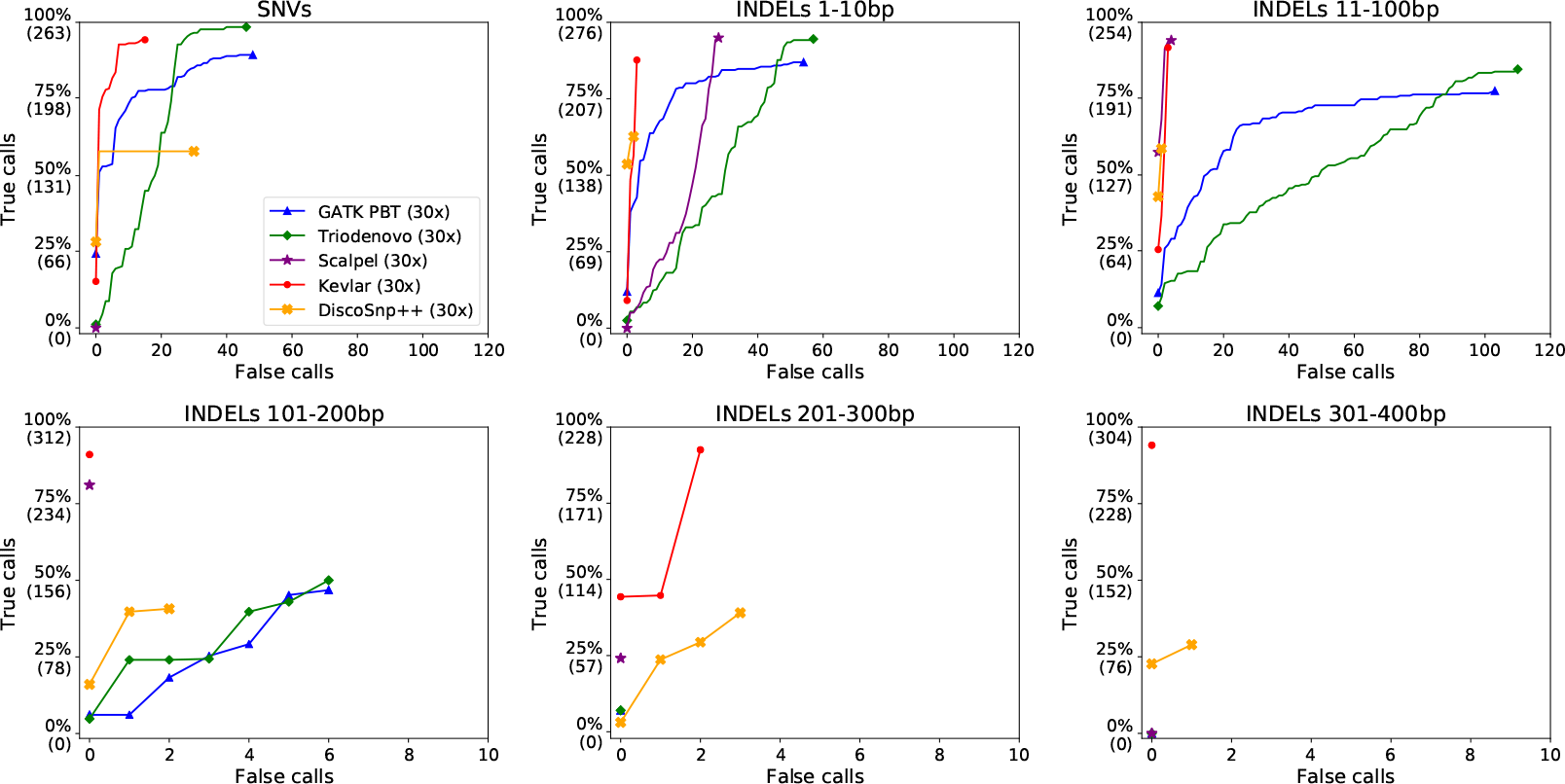
Receiver operating characteristic (ROC) curves comparing variant prediction performance on a simulated data set. Average sequencing depth was approximately 30x. Each of the six panes shows prediction accuracy for a different variant type: single nucleotide variants (SNVs); insertions or deletions (indels) 1-10bp in length; 11-100bp indels; 101-200bp indels; 201-300bp indels; and 301-400bp indels. Note that the scale of the X-axis scale for long indels is an order of magnitude smaller than the X-axis scale for SNVs and short (< 100 bp) indels.

### Performance on the SSC 14153 autism trio

To assess Kevlar’s performance on real data, we applied Kevlar to predict *de novo* variants in the proband of an autism trio from the Simons Simplex Collection (family 14153). As a reference for comparison, we obtained a potential “truth set” from the denovo-db database (http://denovo-db.gs.washington.edu/denovo-db/). This truth set includes 196 *de novo* variant predictions, and represents the union of predictions made for this trio by several recent studies.^39–41^ Note that the expected number of *de novo* variants per generation is estimated to be around 100,^17, 40^ or about half of the number of predictions in the truth set. Annotations in the denovo-db database indicate that experimental validation has confirmed 14 of the 196 calls.

In total, Kevlar predicts 219 *de novo* variants for trio 14153, including 150 SNVs, 68 indels/SVs, and a single 2 bp multinucleotide variant (MNV). We note that Kevlar assigned many of these predicted variants a low likelihood of the variant being a true *de novo* event. Figure 3 shows the congruence between the 100 top ranked Kevlar calls and the denovo-db calls for this trio.

**Fig 3.**
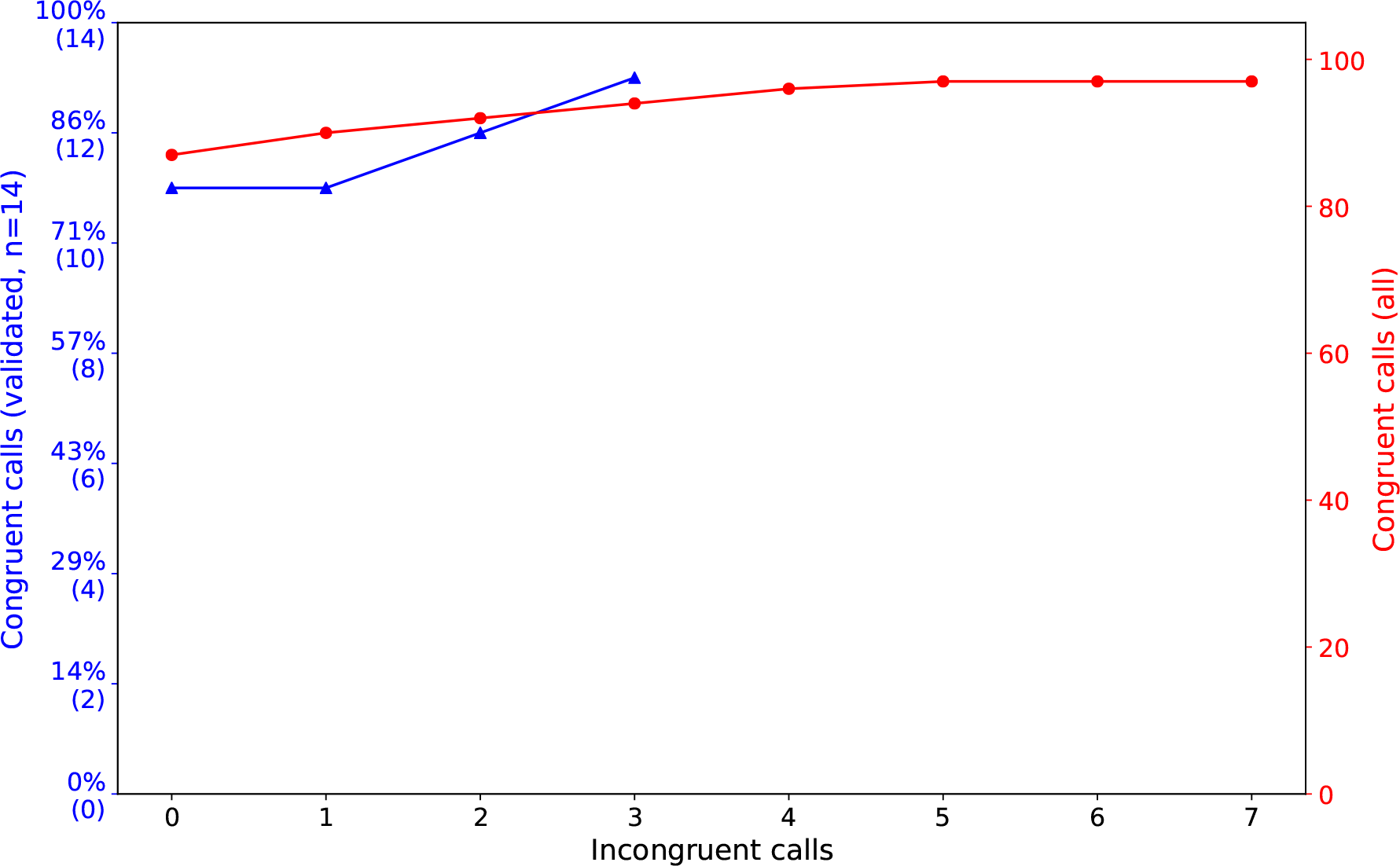
ROC curves showing congruence between *de novo* variant calls made by Kevlar on the SSC 14153 trio and corresponding calls from the denovo-db variant database. The red curve shows Kevlar’s performance compared to all denovo-db calls, and the blue curve shows Kevlar’s performance compared to denovo-db calls with experimental validation.

Of the 14 denovo-db calls with experimental validation, 13 (92.9%) were predicted accurately by Kevlar and assigned a high likelihood score indicative of a confident *de novo* variant call. Overall, the 100 Kevlar variant calls ranked highest by the likelihood score include only 4 calls not present in denovo-db (probable false calls). On the other hand, only 5 Kevlar variant calls present in denovo-db (probable true variants) are not among the 100 highest ranked Kevlar calls. Of the 196 denovo-db calls, 95 are absent from the Kevlar predictions. The majority of these calls (75 / 95, ≈80%) occur in regions of repetitive DNA and have shown to be unreliable in experimental validation (Tychele Turner, personal communication).

Finally, a recent study verified the presence of a *de novo* deletion of approximately 6kbp in the proband of this trio,^40^ removing the 5’UTR of the gene *CANX*. Kevlar also predicted this *de novo* deletion successfully, and identified the precise (and previously undetermined) breakpoints at chr5:179,122,593 and chr15:179,128,130 (GRCh37). Inspection of the variant reveals that both of the deletion’s breakpoints occur in *Alu* repeat elements abundant throughout the genome (Figure 4). As a result, only 7 of the *k*-mers spanning the variant are unique signatures of mutation not already present elsewhere in the genome. We observe with interest that both breakpoints occur inside a 20 bp identical repeat, indicating this *de novo* deletion is the result of non-allelic homologous recombination (NAHR).

**Fig 4.**
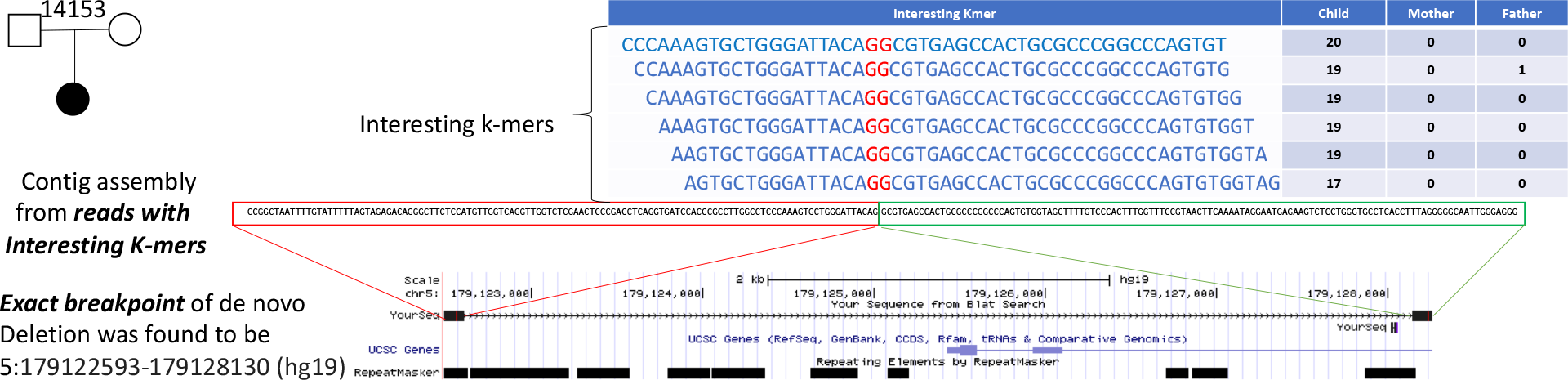
The validated 6kbp *de novo* deletion as predicted by Kevlar. The interesting *k*-mers, their abundances in each sample, the assembled novel sequence, and the breakpoint are depicted.

## Discussion

*De novo* variants are a major contributing factor in many disorders (e.g. intellectual disability, autism, and epilepsy). Accurate discovery of these variants has been challenging as prediction methods need to be confident not only in the existence of the event in the proband/child, but also in the absence of the variant in the parents. Current approaches depend on correct alignments of sequence reads to a reference genome. Any complications in computing read alignments due to repeats, gaps in the reference, or variant complexity can result in false predictions or failure to discover a true *de novo* variant.

The method proposed in this study compares *k*-mers between related individuals to find the *k*-mers indicating a *de novo* variant in the sample of interest. We acknowledge the recently proposed methods NovoBreak^30^ and HAWK,^28^ which are conceptually similar and likewise capable of accurately predicting *de novo* variants. Kevlar and related methods do not depend on mapping reads to a reference genome, but instead rely on direct comparison of sequence content between related individuals. This strategy enables Kevlar to accurately predict several classes of *de novo* mutation (substitutions, insertions, deletions, SVs) simultaneously with a single simple mathematical model. As long as the *de novo* mutation creates a *k*-mer not already present in the reference genome, the proposed algorithm should be able to accurately discover the event. We have also developed a *k*-mer based likelihood model for scoring and ranking variant calls according to their probability of being true *de novo* events. This likelihood score is effective in discerning *de novo* variants from inherited mutations and false variant calls. We have demonstrated the effectiveness of our discovery method and scoring model using both simulated and real data. Kevlar is competitive with best-in-class tools for discovery of variety of variant types, and substantially outperforms available methods for discovery of larger *de novo* variants. Not only does Kevlar predict indels and SVs with high sensitivity and specificity, but it reports the exact breakpoints of these variants with single base pair precision.

Development of completely reference-free methods is tremendously valuable in scenarios where the availability, quality, or relevance of a reference genome is insufficient. Kevlar’s preliminary steps—identifying variant-spanning reads, binning reads into groups corresponding to distinct putative variants, and assembling each read group into a variant-spanning contig—are performed without the use of a reference genome. We note, however, that subsequent steps in the Kevlar workflow to annotate, filter, and score the preliminary variant calls still depend on a reference genome. One promising approach to developing a completely reference-free *de novo* variant discovery method would be to annotate of variants by aligning variant-spanning contigs directly to an assembly/variation graph.

## Methods

### Assessing diagnostic utility of novel k-mers

We expect that a *de novo* mutation will result in numerous novel *k*-mers, given a sufficiently large value of *k*. We also expect that these novel *k*-mers will be present in high abundance, given sufficiently deep sampling of the proband genome. Intuitively, we can use these novel *k*-mers to identify reads that span the *de novo* variant.

We assessed this intuition by traversing the human reference genome (GRCh38) base by base, simulating variants (SNVs and 5 bp deletions) at each position. For each simulated mutation, we determined the fraction of *k*-mers spanning the mutation that exist nowhere else in the genome, and thus act as a diagnostic signature of that particular variant. We then aggregated over the entire genome the probability that *k*-mer spanning a mutation (in this case 31-mers) will be novel—see Figure 1b and 1c.

Based on the results of this experiment, we formulate the *de novo* variant discovery problem as a search for putatively novel *k*-mers that are abundant in the proband and effectively absent in each parent. For sake of simplicity, we are using the term *proband* to refer generally to the subject or focal individual, and *parent* to refer generally to control individuals.

Here, *abundant* and *effectively absent* are defined in terms of a simple threshold model. Let *X* be the *absence threshold*, and *Y* be the *presence threshold*, and *A* = {*A*_*p*_, *A*_*m*_, *A*_*f*_} be the abundances of a *k*-mer in the proband, mother, and father. We designate this *k*-mer as “*interesting*” (putatively novel) if and only if *A*_*p*_ ≥ *Y*, *A*_*m*_ ≤ *X*, and *A*_*f*_ ≤ *X*. Based on our experience, the values *Y* = 5 and *X* = 1 produce desirable results for 30x sequencing coverage.

### Kevlar workflow

The steps of the Kevlar workflow, summarized at a high level in Figure 1d, are described in detail in the subsequent sections.

### Step 0: Compute k-mer counts

Preliminary to identifying novel *k*-mers, the abundance of each *k*-mer in each sample must be counted. Storing exact counts of every *k*-mer requires a substantial amount of space (dozens of gigabytes or more per sample), so Kevlar exploits several strategies to reduce the space required for keeping *k*-mer counts in memory.

First, Kevlar stores approximate *k*-mer counts in a Count-Min sketch, a probabilistic data structure similar to a Bloom filter that operates in a fixed amount of memory, exchanging accuracy for space efficiency.^42^ The accuracy of the Count-Min sketch depends on its size and the number of distinct elements (*k*-mers in this case) being tracked. The Count-Min sketch exhibits a one-sided error, meaning that *k*-mer counts are sometimes overestimated but never underestimated. The extent of inaccuracy in the *k*-mer counts is summarized by the *false discovery rate (FDR)* statistic computed from the occupancy of the Count-Min sketch.

Second, Kevlar uses a masked counting strategy in which *k*-mers present in the reference genome and a contaminant database are ignored. This substantially reduces the number of *k*-mers to be stored in the Count-Min sketch, and as a consequence the desired level of accuracy can be maintained using a smaller amount of space.

Third, *k*-mer counts are recomputed with exact precision in subsequent steps of the Kevlar workflow, which means any *k*-mer retained erroneously due to an inflated count can be compensated for at a later stage. As a consequence, it is possible to reduce the size of the Count-Min sketch even further, resulting in a FDR of 0.5 or greater. (Inicidentally, *k*-mers erroneously *discarded* due to inflated counts cannot be compensated for later, putting an upper bound on the imprecision of the initial *k*-mer counts in the control samples.)

Kevlar’s *k*-mer counting operations are invoked with the `kevlar count` command, and rely on bulk sequence loading procedures and an implementation of the Count-Min sketch data structure from the khmer library.^22, 42, 43^

### Step 1: Identifying novel k-mers and reads

To identify sequences spanning *de novo* variants, Kevlar scans each read sequenced from the proband. The per-sample abundances of each *k*-mer are queried from the Count-Min sketches computed in previous steps. If a *k*-mer is present in high abundance in the proband and absent from the parents (that is, it satisfies user-specified abundance thresholds), it is designated as “interesting” or putatively novel. This operation is similar to the selection of “novo” *k*-mers by NovoBreak^30^ and “significant” *k*-mers by HAWK.^28^ Any read containing one or more interesting *k*-mers is retained for subsequent processing. This step is implemented in the `kevlar novel` command.

### Step 2: Contamination, reference, and abundance filters

Reads containing putative novel *k*-mers are filtered prior to subsequent analysis. This filtering step serves two purposes.

First, Kevlar re-computes the abundance of each interesting *k*-mer in the proband sample. The relatively small volume of these reads allows Kevlar to re-compute *k*-mer counts with perfect accuracy in a small amount of memory and time. Any *k*-mer whose corrected count no longer satisfies the required abundance threshold is discarded. Note that since only proband reads are retained, only the proband *k*-mer abundances can be recomputed. This filtering step will not recover a *k*-mer that is erroneously discarded in the previous step due to an erroneously inflated *k*-mer abundance in one of the control (parent and sibling) samples.

Second, if for any reason *k*-mers from the reference genome and contaminants are not ignored in the initial *k*-mer counting step, this filtering step provides another opportunity to discard these *k*-mers.

After these filters are applied, any read that no longer contains any novel *k*-mers is discarded, and the remainder of the reads are retained for subsequent analysis.

The `kevlar filter` command is used to execute these contamination, reference, and abundance filters.

### Step 3: Partitioning reads using shared novel k-mers

Interesting reads spanning the same variant are expected to share numerous interesting *k*-mers. These shared novel *k*-mers provide a mechanism for grouping the reads into disjoint sets reflecting distinct variants.

To be precise, we define a *read graph G* as follows: every read containing one or more novel *k*-mers is represented by a node in *G*, and a pair of nodes is connected by an edge if they have one or more novel *k*-mers in common. With this formulation, if two reads share a novel *k*-mer they must be part of the same connected component in *G*. Overall *G* is sparse, but typically each connected component of the graph is highly connected. In subsequent steps, each component or partition *p ∈ G* is analyzed independently.

The `kevlar partition` command implements this partitioning strategy.

### Step 4: Contig assembly and reference target selection

For each connected component *p* ∈ *G*, we assemble the corresponding reads using the overlap-based algorithm implemented in the fermi-lite library.^44^ Briefly, fermi-lite performs error correction, trims reads at unique *l*-mers, constructs an FM-index of the trimmed reads, and constructs a transitively reduced overlap graph. The optimal path in the final graph is output as a contig *C*_*p*_ suitable for variant calling.

Next, we select a target reference sequence (or set of candidate targets) for the contig *C*_*p*_. Briefly, Kevlar decomposes the contig into overlapping subsequences of length *l*(*seeds*; *l* = 51 by default), and uses BWA MEM^45^ to identify locations of exact matches for each seed sequence in the reference genome. The genomic interval that spans all seed exact matches, plus Δ nucleotides in each direction (Δ = 50 by default), is then selected as the target reference sequence for *C*_*p*_. If any adjacent seed matches are separated by more than *D* nucleotides (*D* = 10,000 by default), then the seed matches are split at that point and multiple reference targets are selected. The set of reference target sequences corresponding to contig *C*_*p*_ is denoted 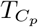.

Read assembly is invoked with the `kevlar assemble` command, and reference target selection is invoked with the `kevlar localize` command.

### Step 5: Contig alignment and variant annotation

The contig *C*_*p*_ is aligned to each reference target sequence 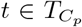 using the ksw2 library^46^—specifically its implementation of Green’s formulation of dynamic programming global alignment and extension (`ksw2 extz`). If there are multiple candidate targets, only the highest scoring alignment is retained. When a contig aligns to multiple locations with the same optimal score, all optimal alignments are retained for variant calling.

Prior to variant calling, kevlar right-aligns any gaps at the right end of the alignment to minimize the number of alignment blocks/operations. Next, Kevlar inspects the alignment path (represented as a CIGAR string) of each alignment and tests for matches against expected patterns. Alignments matching the pattern `^(\d+[DI])?\d+M(\d+[DI])?$` are classified as SNV events, and the “match” block of the alignment is scanned for mismatches between the contig and the reference target. Any mismatch is reported as a single nucleotide variant. Alignments matching the pattern `^(\d+[DI])?\d+M\d+[ID]\d+M(\d+[DI])?$` are classified as indel events. In addition to reporting the internal gap of this alignment as an indel variant, the flanking “match” blocks are also scanned for mismatches between the contig and the target to be reported as putative SNVs. Any alignment not matching the two patterns described above is designated as an uninterpretable “no-call” and listed in the output along with the corresponding contig sequence.

In some cases, there is a possibility that kevlar will report two or more calls in close proximity. While the probability of two *de novo* variants occurring in close proximity is effectively nil, it is common for an inherited variant to occur proximal to a *de novo* variant. Occasionally one of these inherited variants will not be spanned by any interesting *k*-mers, in which case it can immediately be designated as a “passenger” variant call. However, in cases where an inherited variant is spanned by one or more interesting *k*-mers, we rely on subsequent examination of *k*-mer abundances to distinguish novel variants from inherited variants.

The `kevlar call` command computes the contig alignments and makes preliminary variant calls.

### Step 6: Likelihood scoring model for ranking and filtering variant calls

Given the filters already discussed, false *interesting k*-mer designations are rare throughout the genome overall. Redundancy from a high depth of sequencing coverage prevents sequencing errors from driving the reported abundance of *k*-mers present in the parents to 0. And for loci where local coverage is low in one parent, it is extremely rare that local coverage will also be low in the other parent by chance. Thus, if a *k*-mer is present in either parent, it is disqualified from the *interesting* or novel designation.

However, false *interesting k*-mer designations are enriched around inherited mutations. It is very common for variants present in one parent to be absent from the other parent. If by chance the depth of sequencing coverage is low at such a locus in the donor parent, there may not be enough redundancy to compensate for sequencing errors. As a result, some *k*-mers that are truly present in the donor parent will have a reported abundance of 0. Being truly absent from the other parent, these *k*-mers are erroneously designated as unique to the proband.

A related complication occurs when a novel variant is proximal to an inherited variant. Both variants are reflected in the alignment of the associated contig (assembled from proband-derived *interesting* reads) to the reference genome. In both of these cases, distinguishing novel variants from inherited variants benefits from examination of the abundances of all *k*-mers containing each variant, as well as the corresponding reference *k-mers*.

We utilize a likelihood based model to score and rank the predicted *de novo* variants. We consider the abundance of the interesting *k*-mers to calculate the likelihood of the event observed being *de novo*, inherited, or a false call. Using these likelihood probabilities, we calculate a score for each variant being truly a *de novo* variant based on ratio of likelihoods.

First, for each variant we define a set of alternate *k*-mers **A** as the *k*-mers indicating existence of the variant (alternate genotype). We consider only *k*-mers that are unique to this variant (that is, they don’t appear in any other location in the reference genome). We assume that there are a total of *n* alternate *k*-mers.

Let the random variables *v*_*c*_, *v*_*f*_, and *v*_*m*_ indicate the genotype (i.e. {0/0, 0/1, 1/1}) of the putative variant in the proband/child, father, and mother respectively. The random variable 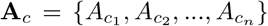 denotes the counts of the alternate allele *k*-mers in the proband, 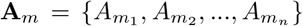 the alternate allele *k*-mer counts in the mother, and 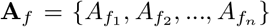 the alternate allele *k*-mer counts in the father. The likelihood that a putative variant is *de novo* can be calculated as follows.

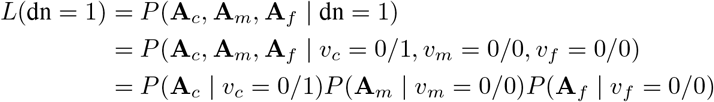

We note that there are dependencies between *k*-mer counts within a sample. However, to simplify the calculation of likelihoods we assume independence between *k*-mer counts and provide an approximation of the likelihoods. For calculating the probability of an observed *k*-mer count conditioned on a 1/1 genotype, we assume a normal distribution with mean *µ* and standard deviation *σ*. The *µ* and *σ* parameters are learned empirically for each sample using only exonic *k*-mers that occur only once in the reference genome. The probability of an observed *k*-mer count conditioned on a 0/1 genotype is calculated assuming a normal distribution with mean 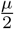 and standard deviation 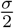. For the genotype 0/0 we use binomial distribution to calculate the likelihood of the observed *k*-mer abundance assuming the *k*-mer is generated by, e.g., sequencing error. Similarly, we calculate the likelihood that a putative variant is a false positive prediction by conditioning on the variant’s non-existence (genotype 0*/*0) in all three samples, i.e. *L*(*f*_*p*_ = 1) = *P*(***A***_*c*_, ***A***_*m*_, ***A***_*f*_ | *v*_*c*_ = 0/0,*v*_*m*_ = 0/0,*v*_*f*_ = 0/0).

Finally, we calculate the likelihood of observed *k*-mer counts under the inheritance assumption. As there are several different valid scenarios to represent variant inheritance the likelihood calculation requires additional steps as explained below (again assuming independence of *k*-mer abundances as an approximation).

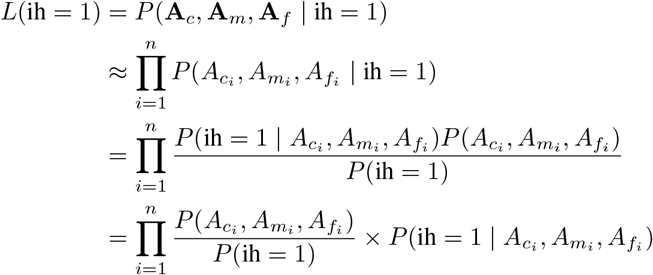

We calculate the 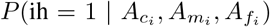 as summation of probability of possible trio-genotype combinations representing inheritance scenarios (e.g., (*v*_*c*_ = 1/0, *v*_*f*_ = 1/0, *v*_*m*_ = 0/0) or (*v*_*c*_ = 1/0, *v*_*f*_ = 0/0, *v*_*m*_ = 1/0)). Furthermore, we assume a constant prior value for *P*(ih = 1) based on all possible valid inheritance scenarios.

Finally, we utilize a score motivated from the likelihood ratio test to score and rank any predicted variant as being a *de novo* variant. Note that, as numerical calculation of the likelihoods is numerically prone to error we consider the logarithm of the score. Thus, we formally define the score assigned to each variant for being *de novo* as *S*_*L*_ = log*L*(*dn* = 1) − max{log(*L*(*ih* = 1))}, log(*L*(*f*_*p*_ = 1)). The `kevlar simlike` command computes likelihoods for preliminary variant calls, sorts the calls, and filters out low scoring and otherwise problematic calls.

### Data simulations

We simulated whole-genome shotgun sequencing for a hypothetical trio (father, mother, and proband) to evaluate the accuracy of our *de novo* variant discovery algorithm. Using the human reference genome (GRCh38) and a catalog of common variants (dbSNP), we constructed two independent diploid genomes representing the two parents. We randomly selected SNPs and indels from dbSNP and assigned the variants to each parental haplotype at a rate of 1 for every 1000 bp.

We then constructed the diploid proband genome through recombination of the parental diploid genomes and simulated germline mutation. SNVs and indels ranging from <10 bp to 400 bp in length were simulated as heterozygous events unique to the proband, representing *de novo* variation.

Finally, we used wgsim^47^ to simulate whole-genome shotgun sequencing of each individual. This produced se-quences resembling Illumina 2×150bp paired-end reads with low sequencing error rate. The sequencing was repeated at four different average depths of sequencing coverage: 10x, 20x, 30x, and 50x.

## Acknowledgements

We would like to acknowledge Dr. Tamer Mansour, Luiz Irber Jr., Camille Scott, and Lisa Johnson for helpful discussions on method development and implementation, and Dr. Tychele Turner for helpful discussions on the method evaluation.

## Author Contributions

DSS, CTB, and FH conceived the study. DSS implemented the method. DSS and FH designed the experiments and wrote the manuscript. DSS, CTB, and FH edited and approved the final manuscript.

## Declaration of Interests

The authors declare no competing interests.

## Appendix

**Fig A1.**
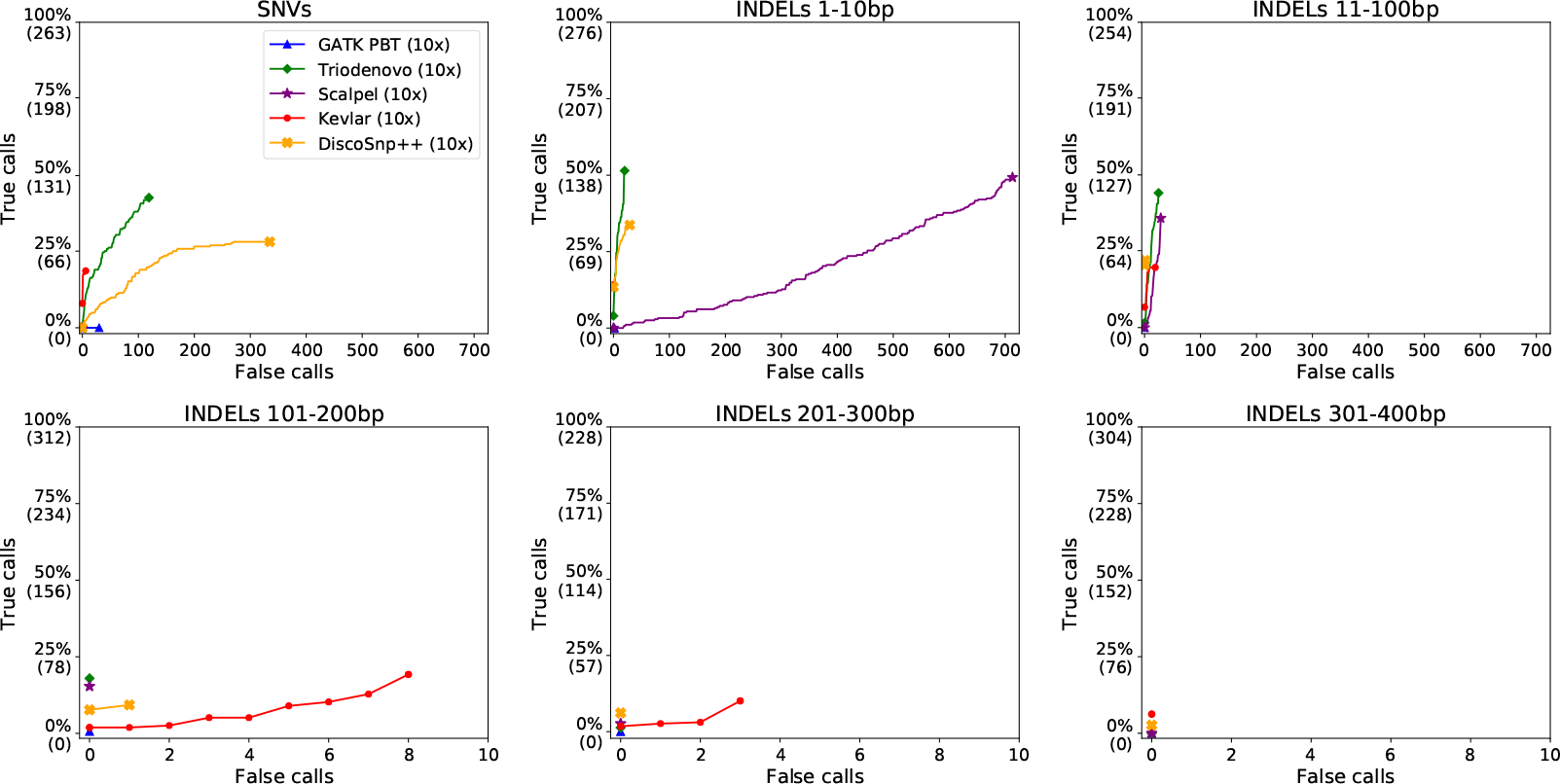
Receiver operating characteristic (ROC) curves comparing variant prediction performance on a simulated data set. Average sequencing depth was approximately 10x. Each of the six panes shows prediction accuracy for a different variant type: single nucleotide variants (SNVs); insertions or deletions (indels) 1-10bp in length; 11-100bp indels; 101-200bp indels; 201-300bp indels; and 301-400bp indels. Note that the scale of the X-axis scale for long indels is an order of magnitude smaller than the X-axis scale for SNVs and short (< 100 bp) indels.

**Fig A2.**
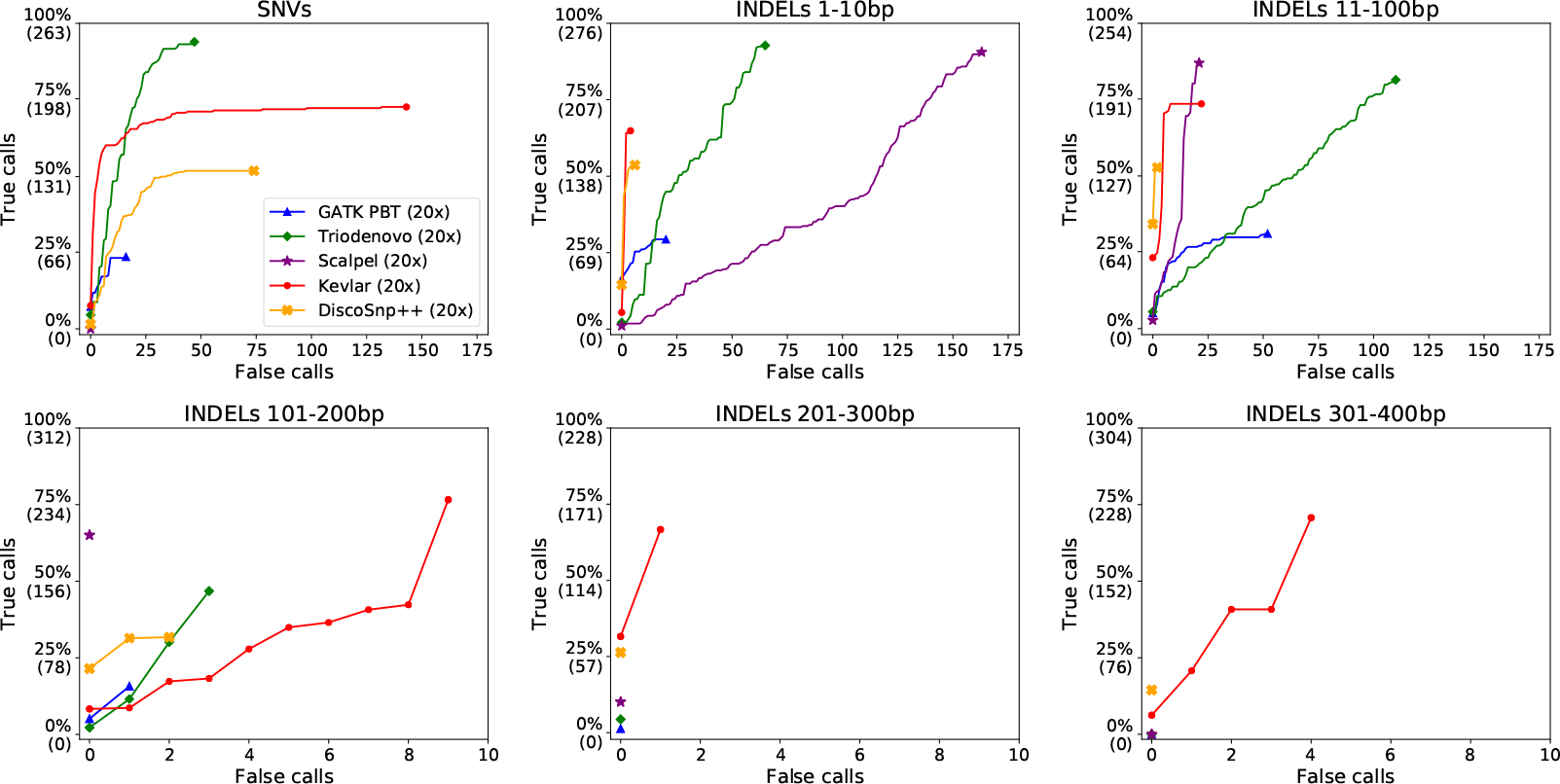
Receiver operating characteristic (ROC) curves comparing variant prediction performance on a simulated data set. Average sequencing depth was approximately 20x.

**Fig A3.**
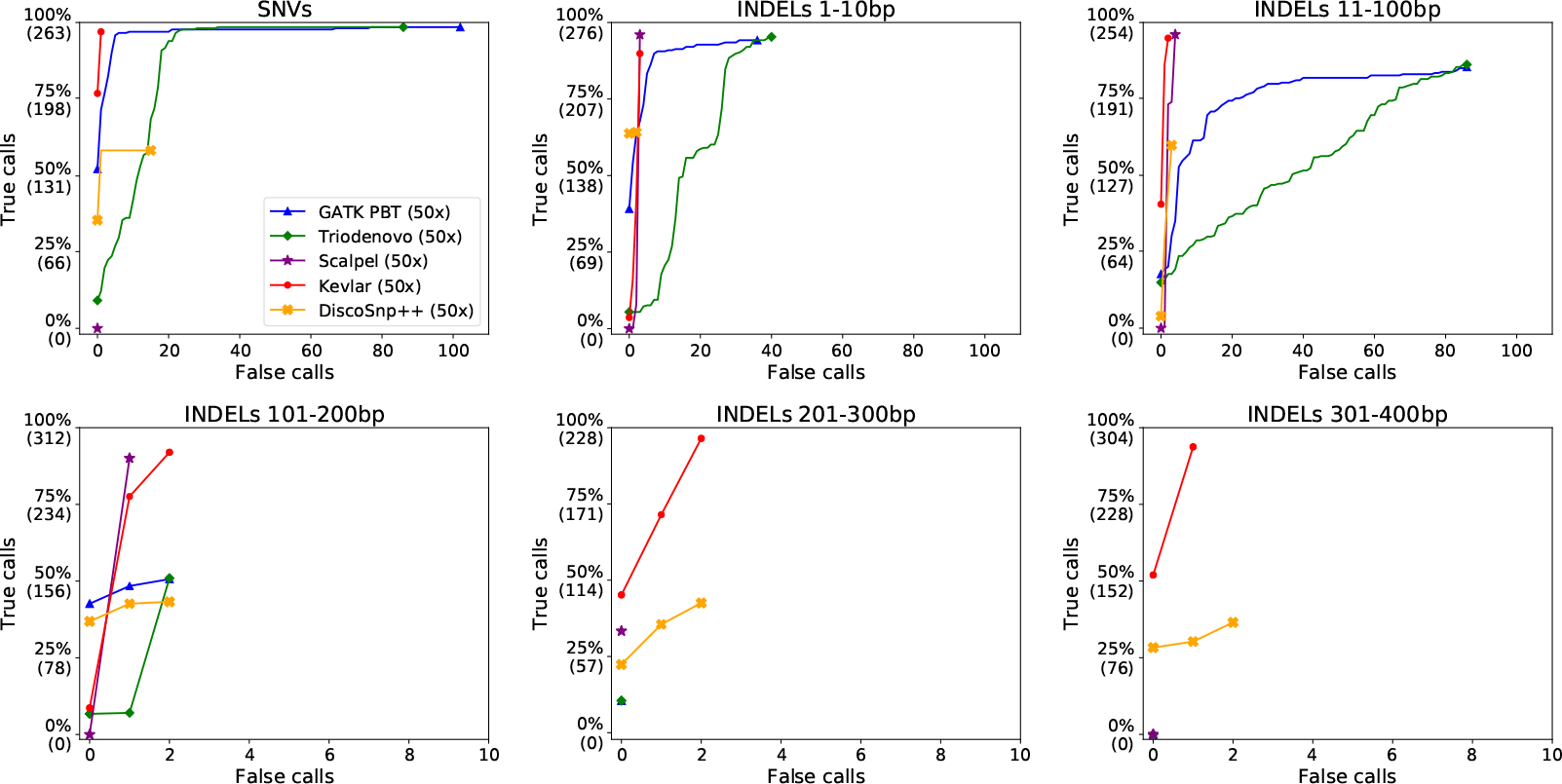
Receiver operating characteristic (ROC) curves comparing variant prediction performance on a simulated data set. Average sequencing depth was approximately 50x.

